# Serial passaging affects stromal cell mechanosensitivity on hyaluronic acid hydrogels

**DOI:** 10.1101/2023.03.16.532853

**Authors:** Jenna L. Sumey, Abigail M. Harrell, Peyton C. Johnston, Steven R. Caliari

## Abstract

There is tremendous interest in developing hydrogels as tunable *in vitro* cell culture platforms to study cell response to mechanical cues in a controlled manner. However, little is known about how common cell culture techniques, such as serial expansion on tissue culture plastic, affect subsequent cell behavior when cultured on hydrogels. In this work we leverage a methacrylated hyaluronic acid hydrogel platform to study stromal cell mechanotransduction. Hydrogels are first formed through thiol-Michael addition to model normal soft tissue (e.g., lung) stiffness (*E* ~ 1 kPa). Secondary crosslinking via radical photopolymerization of unconsumed methacrylates allows matching of early- (*E* ~ 6 kPa) and late-stage fibrotic tissue (*E* ~ 50 kPa). Early passage (P1) primary human mesenchymal stromal cells (hMSCs) display increased spreading, myocardin-related transcription factor-A (MRTF-A) nuclear localization, and focal adhesion size with increasing hydrogel stiffness. However, late passage (P5) hMSCs show reduced sensitivity to substrate mechanics with lower MRTF-A nuclear translocation and smaller focal adhesions on stiffer hydrogels compared to early passage hMSCs. Similar trends are observed in an immortalized human lung fibroblast line. Overall, this work highlights the implications of standard cell culture practices on investigating cell response to mechanical signals using *in vitro* hydrogel models.

## 1. Introduction

Mechanotransduction, or the process by which cells sense and interpret biomechanical cues from their environment^1^, is a major contributor in tissue homeostasis as well as during development, injury response, and pathological disorders^2^. Of particular interest are cell-microenvironment interactions during aberrant tissue remodeling processes like fibrosis, a gradual stiffening process characterized by excessive scarring that results in decreased organ compliance and eventual failure^3^. Fibrosis, which can affect many organs including the heart, liver, and lungs^4^, occurs in part due to positive feedback loops in the injury response cascade. Activated cells (myofibroblasts) deposit excessive extracellular matrix (ECM) and lead to gradual tissue stiffening, which serves as a mechanical stimulus to perpetuate myofibroblast activation^5^. Studies have shown that elevated substrate stiffness alone drives activation of quiescent cells into pro-fibrotic myofibroblasts in both fibrosis^6–10^ and cancer^11,12^. However, understanding the specific role of biomechanical cues on cellular activation is complicated in the native tissue microenvironment due to the presence of many interrelated biochemical and biophysical cues.

To address the challenges associated with studying cell behavior *in vivo*, hydrogels have recently become a popular class of biomaterials for *in vitro* cell culture applications since they can be designed to mimic relevant aspects of the native ECM, including mechanical properties such as Young’s modulus or stiffness in the range of kilopascals (kPa)^13^, as opposed to traditional tissue culture plastic (TCP) that is supraphysiologically stiff^14^. Hydrogels can be engineered to match physiologic ranges of tissue mechanics by tuning parameters including the polymer concentration, crosslinking density, and crosslinking type (e.g., physical and/or covalent). Hydrogel systems incorporating mechanical cues have been used in evaluating cell spreading and cytoskeletal organization^15,16^, stem cell differentiation^17–19^, and myofibroblast behavior in heart^20^, lung^21,22^, and liver^23^ disease. Stiffer substrate mechanics have been shown to support focal adhesion (FA) formation, which includes proteins like paxillin, and assist in regulating changes in cell behavior such as spreading and differentiation^24–28^. As FAs mature and grow from < 0.25 μm to 1-5 μm^29–31^, they facilitate cytoskeletal polymerization of actin stress fibers^30,32^ following the nuclear localization of myocardin-related transcription factor-A (MRTF-A)^33^, which is involved in pro-fibrotic gene expression^34–38^.

Understanding cell response to mechanical cues is especially important given recent evidence suggesting that cells maintain a ‘memory’ of their surrounding mechanical environment, resulting in changes in cell sensitivity to matrices with different mechanical cues^39–46^. Seminal reports on mechanical memory showed that fibroblasts cultured extensively on stiffer silicone-based biomaterials maintained an activated myofibroblast phenotype, even when moved to a softer substrate^40^. Conversely, fibroblasts primed on softer substrates showed blunted activation when moved to stiffer substrates^40^. Subsequent work showed that mechanical memory was regulated in part by MRTF-A, a transcriptional regulator implicated in the upregulation of profibrotic genes such as *Acta2^44^*.

Collectively, these studies suggest that the extent of mechanical dosing plays a key role in mechanotransduction signaling and cell phenotype. However, it is not well understood how standard culture techniques, such as serial passaging on TCP, influence resultant cell behavior in hydrogel cell culture models. In this study, we compared the response of both primary human mesenchymal stromal cells (hMSCs), which can play a role in multiple fibrotic pathologies, as well as immortalized human lung fibroblasts to engineered hydrogels that matched the stiffness of either normal or diseased soft tissue to determine how mechanosensitivity changed with respect to initial culture and expansion on TCP. Overall, these results describe how the mechanical sensitivity of hMSCs and fibroblasts in experimental hydrogel models of fibrosis changes with prior expansion on TCP.

## 2. Materials and Methods

### 2.1 MeHA synthesis

Hyaluronic acid (HA) was methacrylated as previously described^23^. Sodium hyaluronate (Lifecore, 60 kDa) was dissolved at 2 wt% in deionized water prior to reacting with methacrylic anhydride (Sigma Aldrich, 4.83 mL per g HA) at pH 8.5-9 for 6 h on ice. After all of the methacrylic anhydride was reacted, the solution was allowed to stir at room temperature overnight. The mixture was dialyzed against deionized water (SpectraPor, 6-8 kDa molecular weight cutoff) at room temperature for 5 days, then frozen and lyophilized until dry. The degree of modification as determined by ^1^H NMR (500 MHz Varian Inova 500) was ~ 100%.

### 2.2 Hydrogel fabrication

MeHA in 0.2 M triethanolamine (TEOA, Sigma Aldrich) buffer at pH 9 was functionalized with a thiolated cell-adhesive RGD peptide (GenScript, GCGYGRGDSPG) via a Michael-type addition reaction. The solution was incubated at room temperature for at least 1 h and afforded a final RGD concentration of 1 mM. 1 kPa MeHA hydrogels were also formed through Michael-type addition. 4 wt% RGD-modified MeHA was crosslinked with dithiothreitol (DTT, Sigma Aldrich) at pH 9. The hydrogel precursor solution (50 μL) was placed between untreated and thiolated glass coverslips (18 x 18 mm) and allowed to crosslink for 1 h at 37°C. Both the RGD functionalization and hydrogel crosslinking for the 1 kPa formulation only consumed ~ 15% of the available methacrylate groups, leaving the remaining available for secondary crosslinking.

### 2.3 Secondary hydrogel stiffening

1 kPa MeHA hydrogels were stiffened prior to cell culture to generate moderate and high stiffness hydrogels. For mechanical testing, 1 kPa hydrogels were incubated in PBS containing 2.2 mM lithium acylphosphinate (LAP) photoinitiator at 37°C for 30 min, then exposed to blue (400-500 nm, 5 mW cm^-2^) light for various amounts of time using an OmniCure S2000 curing light. Following light exposure, hydrogels were rinsed three times with PBS to remove LAP and replaced either with fresh PBS for mechanical testing or media for cell culture.

### 2.4 Mechanical characterization

Initial network formation was tracked through rheology on an Anton Paar MCR 302 rheometer with a cone-plate geometry (25 mm diameter, 0.5°, 25 μm gap) set to 37°C. Hydrogel mechanical properties were assessed at least 24 h after swelling using a displacement-controlled nanoindenter (Optics 11 Piuma). A spherical borosilicate glass probe with a radius of 50 μm and a cantilever stiffness of 0.5 N/m was indented onto the surface of MeHA hydrogels submerged in PBS. The Young’s modulus was determined through the loading portion of the generated force versus distance indentation curve using the Hertzian contact mechanics model and assuming a Poisson’s ratio of 0.5. Each sample was indented 25 times with three replicates per group. Hydrogel topography was mapped using matrix indentations of 5 x 5 grids.

### 2.5 Cell culture

Human bone marrow aspirates (Lonza) were purchased to isolate primary hMSCs. Primary hMSCs were used at passage 1 (P1) or 5 (P5) for experiments. Briefly, bone marrow from a single non-smoking donor was vortexed with ammonium chloride (Stem Cell Technologies) at 200 rcf for 5 min, then placed on ice for 10 min to lyse red blood cells. Cells were washed twice with growth media containing minimum essential medium α (MEM-α, Gibco) supplemented with 16.7 v/v% fetal bovine serum, mesenchymal stem cell-qualified (Gibco), 1 v/v% L-glutamine (Gibco), and 1 v/v% streptomycin/amphotericin B/penicillin at 10,000 μg mL^-1^, 25 μg mL^-1^, and 10,000 units mL^-1^, respectively (Gibco). Human lung fibroblasts (abm hTERT T1015) were used at passage 1 (P1) or 10 (P10) for experiments. Cell culture media contained Dulbecco’s modified Eagle’s medium (DMEM) supplemented with 10 v/v% fetal bovine serum (Gibco) and 1 v/v% streptomycin/amphotericin B/penicillin at 10,000 μg mL^-1^, 25 μg mL^-1^, and 10,000 units mL^-1^, respectively (Gibco). Both cell types were cultured to ~ 80% confluency prior to passaging. Passaging occurred every 4 days for hMSCs, and every 2 days for fibroblasts. When confluent, cells were incubated with 0.5% trypsin-EDTA (Gibco) for 6-8 min. Serum-containing media was added to inactivate trypsin, and the cell solution was centrifuged at 200 rcf for 5 min. The supernatant was removed, and the cell pellet was resuspended with warmed media to ensure a final concentration of 250,000 cells/plate. For cell culture on hydrogels, swollen hydrogels (18 x 18 mm), or glass microscope slides (Fisher Scientific) were placed in non-tissue culture treated 6-well plates and sterilized under germicidal UV light for 2 h, then incubated in culture media for at least 30 min before cell seeding. Cells were trypsinized from culture plates and placed on hydrogels at a density of 2 x 10^3^ cells per hydrogel. For all experiments, culture media was replaced every 2 days.

### 2.6 Cell staining, fluorescence imaging, and quantification

For immunostaining, cells on hydrogels were rinsed with PBS, fixed in 10% buffered formalin for 15 min, permeabilized in 0.1% Triton X-100 for 10 min, then blocked in 3% bovine serum albumin in PBS at room temperature for at least 1 h. For visualizing focal adhesions, cells were fixed using a microtubule stabilization buffer^47^ for 10 min at 37°C prior to blocking. Hydrogels were then incubated with primary antibodies at 4°C overnight. Primary antibodies included MRTF-A (mouse monoclonal anti-Mk11 Abcam ab219981, 1:200) or paxillin (mouse monoclonal anti-paxillin B-2, Santa Crux Biotechnology, sc365379, 1:500) to visualize focal adhesions. The following day, hydrogels were washed with PBS three times, then incubated at room temperature in the dark for 2 h with secondary antibodies (AlexaFluor 488 goat anti-mouse IgG 1:400 or 1:600) or rhodamine phalloidin (Invitrogen, R415, 1:600) to visualize F-actin. The hydrogels were rinsed with PBS three more times before incubating with a DAPI nuclear stain (Invitrogen D1306, 1:10,000) for 1 min. The hydrogels were rinsed twice more with PBS and stored in the dark at 4°C prior to imaging. For cell shape and focal adhesion imaging, images were obtained using a Zeiss AxioObserver 7 inverted microscope at 40x oil objective (numerical aperture: 1.3). For quantification of cell spread area, cell shape index (CSI) and MRTF-A nuclear/cytosolic ratio, a CellProfiler (Broad Institute, Harvard/MIT) pipeline was used. CSI quantifies the circularity of the cell, where a line and a circle is characterized by values of 0 and 1 respectively, and was calculated using the formula:

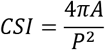

where *A* is the cell area and *P* is the cell perimeter. For focal adhesion analysis, adhesion count and length were quantified using the Focal Adhesion Analysis Server (FAAS)^48^ image processing pipeline with a threshold of 4.5 and a minimum pixel size of 25. At least 20 images were taken of each hydrogel (60 total images per experimental group) for cell shape, MRTF-A, and focal adhesion analyses.

### 2.7 Statistical analysis

For mechanical characterization, at least three hydrogel replicates were used with the data presented as the mean ± standard deviation for hydrogel mechanics, cell shape, and focal adhesion results, and mean ± standard error of mean for CSI and MRTF-A nuclear/cytosol ratio data. One- or two-way analysis of variance (ANOVA) followed by Tukey’s post-hoc tests were performed for all quantitative tests. All cell experiments included at least three replicate hydrogels per group. Box plots of single cell data include mean and median indicators and contain error bars that are the lower result of 1.5 * interquartile range or the maximum/minimum value. Data points between the 1.5 * interquartile range and the maximum/minimum are indicated with circles. Error is reported in single cell figures as the standard error of the mean unless otherwise noted. Significance was indicated by *, **, ***, or **** corresponding to *P* < 0.05, 0.01, 0.001, or 0.0001, respectively.

## 3. Results

### 3.1 Combined Michael-type addition and light-mediated crosslinking chemistries enable the formation of hydrogels matching normal and fibrotic tissue stiffness

Methacrylates were functionalized to the carboxylic acid site on hyaluronic acid to produce hydrogels with sequential crosslinking capabilities (**Fig. S1**)^17,23^. Hydrogels were initially produced through a base-catalyzed Michael-type addition reaction in the presence of dithiothreitol (DTT) to form dithiol crosslinks. This reaction was allowed to proceed for 1 h at 37°C, at which point the storage modulus reached a plateau (**Fig. S2A**). Importantly, during this crosslinking reaction < 15% of the available methacrylates were consumed, leaving the remaining available for secondary crosslinking. The unreacted methacrylates underwent visible light-mediated radical polymerization in the presence of 2.2 mM lithium acylphosphinate (LAP) photoinitiator using blue light (400-500 nm, 5 mW cm^-2^) exposure, creating kinetic chains between the methacrylates (**Fig. 1A**). Adjusting the length of light exposure produced hydrogels with increasing Young’s moduli, as measured by nanoindentation (**Fig. 1B, S2B**). Using these coupled chemistries, hydrogels of 1, 6, and 50 kPa were formed which correlate with the typical Young’s moduli of normal and increasingly fibrotic lung tissue^49^. Hydrogel surface topography and mechanics were relatively homogeneous as shown by spatial mapping obtained via nanoindentation (**Fig. S2C**).

**Figure 1.**
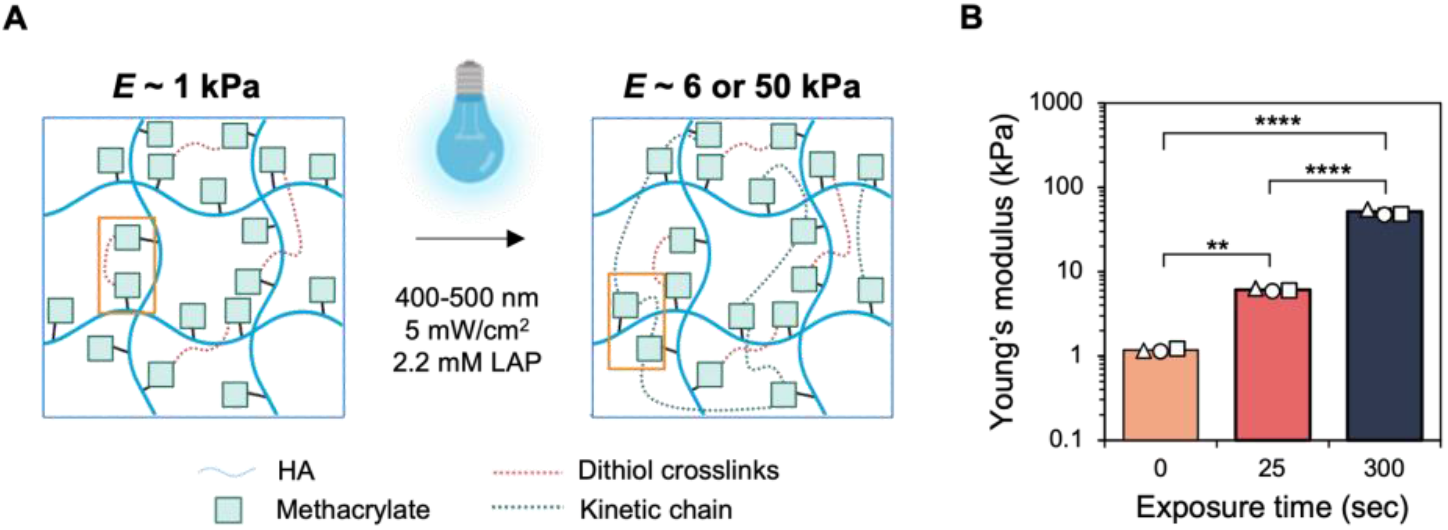
A) Schematic of initial Michael-type addition crosslinking of methacrylates with dithiols to yield a compliant (E ~ 1 kPa) hydrogel, followed by light-mediated radical crosslinking of unconsumed methacrylates to stiffen the substrate. Consuming < 15% of the available methacrylate groups during the initial Michael-type addition allowed for the remaining methacrylates to crosslink upon light exposure. B) Nanoindentation of hydrogels following initial hydrogel formation (1 kPa) and light exposure doses of 25 or 300 sec, yielding hydrogels with Young’s moduli of 6 and 50 kPa respectively. **** P < 0.0001, ** P < 0.01. Data are from n = 3 hydrogel replicates containing at least 75 indentations per group. Data are reported as the mean ± s.d. with the three scatter points on each bar indicating the averages for each hydrogel replicate.

### 3.2 Early passage stromal cells show distinct morphologies as a function of hydrogel stiffness

After characterizing our hydrogel platform, we next seeded early passage stromal cells (either primary human mesenchymal stromal cells (hMSCs) or immortalized human lung fibroblasts) that were grown to confluency on TCP only once (P1) onto MeHA hydrogels of either 1, 6, or 50 kPa. We also seeded cells on glass coverslips (~ GPa) as a control accounting for conventional culture conditions. All cultures were carried out for 4 days. Visual inspection of cells indicated morphological differences across each substrate group, with increased spreading and the formation of organized F-actin stress fibers observed on the 50 kPa and glass matrices (**Fig. 2A**, **S3A**). Quantification of shape metrics showed significant differences in cell spread area as well as cell shape index (CSI), a measure of cell circularity, for the different substrate groups. hMSCs on the 1 and 6 kPa hydrogels displayed similar spreading, while cells on the 50 kPa hydrogel and glass showed no distinct differences in spread area between each other; however, hMSCs on the two softer hydrogel groups showed significantly reduced spread area compared to the two stiffer groups (**Fig. 2B**). Similar trends were observed for fibroblasts (**Fig. S3B**). hMSCs on the 1 kPa hydrogel were significantly rounder (higher CSI) than hMSCs on other substrates (**Fig. 2C**) while fibroblasts were significantly rounder on both 1 and 6 kPa hydrogels compared to stiffer substrates (**Fig. S3C**). The MRTF-A nuclear localization trends were not as distinct for the different groups, with cells displaying significantly higher nuclear translocation on the 50 kPa hydrogel (hMSCs, **Fig. 2D**) or glass substrates (fibroblasts, **Fig. S3D**).

**Figure 2.**
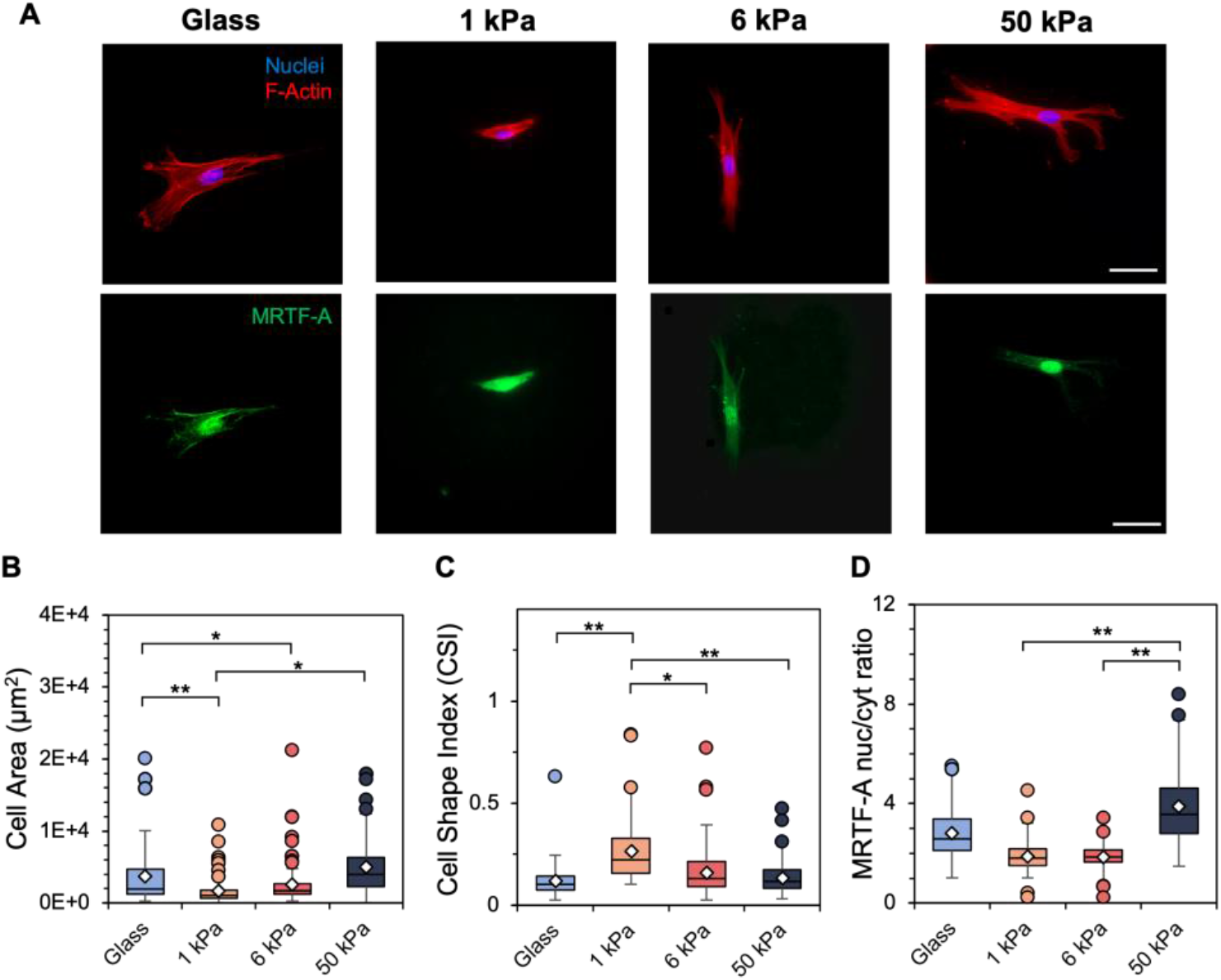
A) Representative images of passage 1 (P1) hMSCs cultured on glass and 1, 6, and 50 kPa hydrogels for 4 days. Scale bars: 50 μm. Following 4 days of culture, quantification of B) cell spread area (μm^2^), C) cell shape index, a measure of cell circularity, and D) MRTF-A nuclear-to-cytosol intensity ratio was performed. *n* = 3 hydrogels per group. ** *P* < 0.01, * *P* < 0.05.

### 3.3 Early passage stromal cells display reduced focal adhesion size on 1 kPa hydrogels

Focal adhesion organization in early passage stromal cells was visualized through paxillin staining, with diffuse staining observed for both hMSCs and fibroblasts seeded on 1 kPa hydrogels, compared to punctate development on all other experimental groups (**Fig. 3A, S4A**). Stromal cells cultured on 1 kPa hydrogels displayed significantly fewer focal adhesions greater than 1.5 μm compared to all other substrate groups (**Fig. 3B, S4B**). Most of the focal adhesions measured ~ 1.3 μm in length for hMSCs and ~ 1.2 μm for lung fibroblasts on 1 kPa hydrogels, while the remaining substrate groups showed a wider range of adhesion lengths with ~ 35-40% of focal adhesions exceeding 1.5 μm (**Fig. 3C, S4C**).

**Figure 3.**
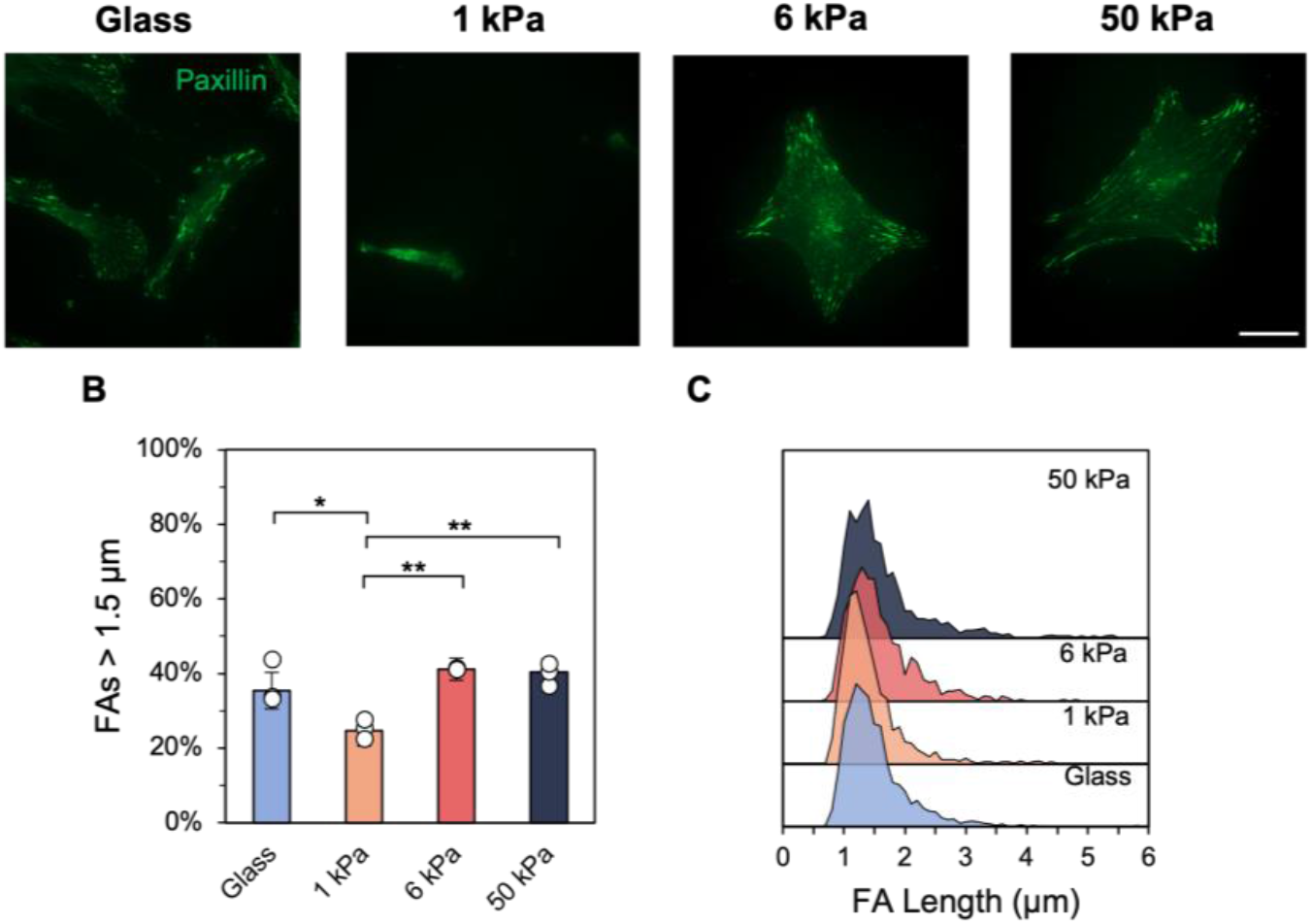
A) P1 hMSCs *were* stained for paxillin to visualize focal adhesions, with quantification of B) focal adhesion lengths larger than 1.5 μm, a metric for mature adhesions, and C) ridgeline plots of the adhesion length distribution. *n* = 3 hydrogels per group. ** *P* < 0.01, * *P* < 0.05.

### 3.4 Late passage hMSCs show similar trends in spreading and roundness compared to early passage hMSCs

Next, to investigate whether serial passaging affected resultant cellular behavior during hydrogel experiments, primary hMSCs were cultured to P5 prior to seeding on hydrogel substrates. Late passage hMSCs largely exhibited similar trends in spreading and roundness as a function of substrate stiffness (**Fig. 4A**) compared to early passage hMSCs. Late passage hMSCs cultured on 1 kPa hydrogels were significantly less spread with lower MRTF-A nuclear localization compared to hMSCs on stiffer substrates (**Fig. 4B, 4D**). Furthermore, late passage hMSCs on compliant hydrogels (1, 6 kPa) were significantly rounder than hMSCs on stiffer surfaces (**Fig. 4C**). Immortalized lung fibroblasts serially passaged to P10 showed more blunted sensitivity to substrate mechanics with significantly lower spreading on 1 kPa hydrogels but similar cell roundness and MRTF-A nuclear localization across all substrate stiffnesses (**Fig. S5**).

**Figure 4.**
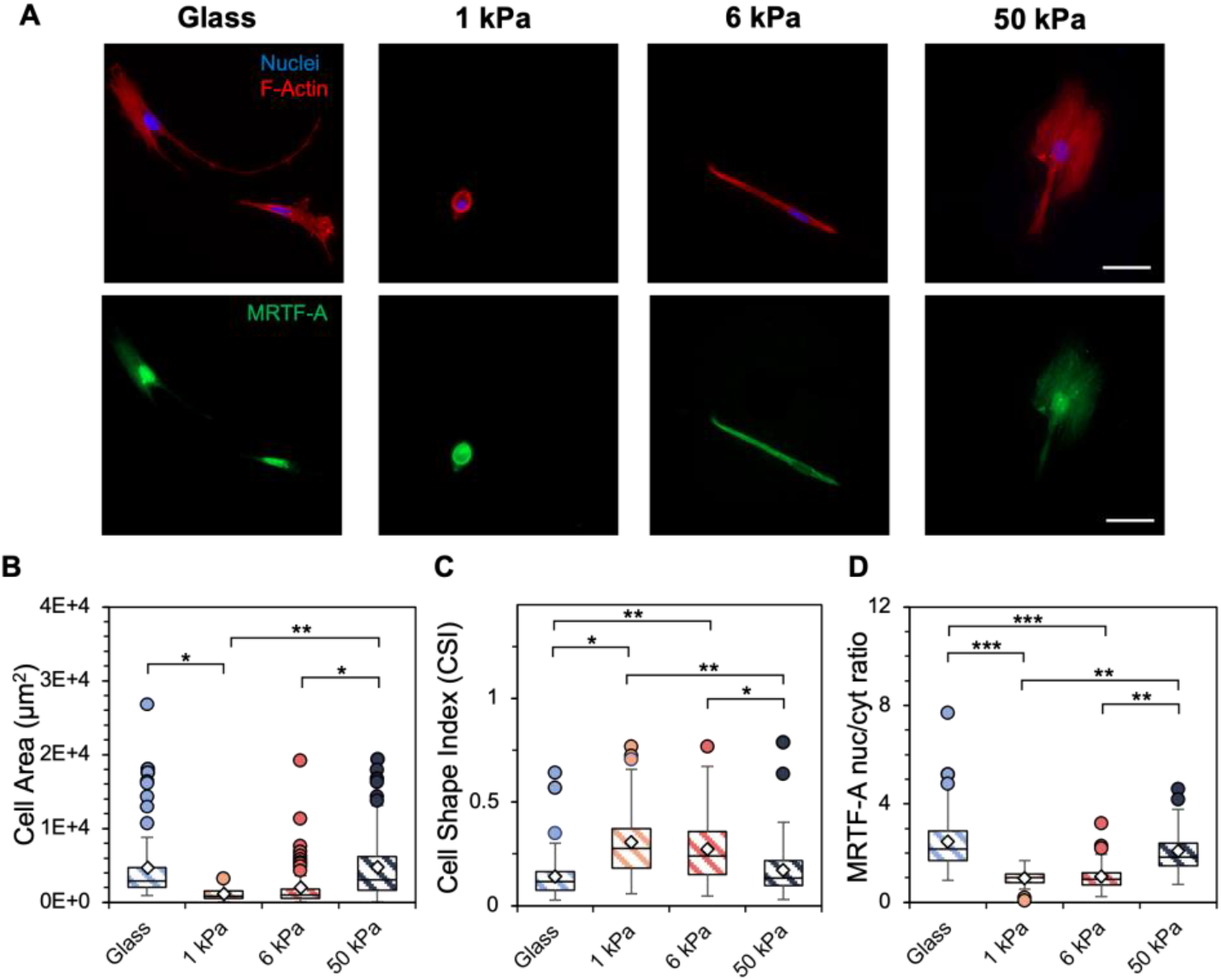
A) Representative images of passage 5 (P5) hMSCs cultured on glass and 1, 6, and 50 kPa hydrogels for 4 days. Scale bar*s*: 50 μm. Following 4 days of culture, quantification of B) cell spread area (μm^2^), C) cell shape index, a measure of cell circularity, and D) MRTF-A nuclear-to-cytosol ratio was performed. *n* = 3 hydrogels per group. *** *P* < 0.001, ** *P* < 0.01, * *P* < 0.05.

### 3.5 Late passage hMSCs show lower MRTF-A nuclear localization and smaller focal adhesions on stiffer hydrogels compared to early passage hMSCs

Late passage stromal cells showed limited differences in focal adhesion organization as a function of substrate stiffness (**Fig. 5**, **S6**). Punctate adhesions are observed for all groups, including some cells on the most compliant 1 kPa hydrogel (**Fig. 5A, S6A**). Similar percentages of mature adhesions above 1.5 μm (~ 30%) were measured for both late passage hMSCs and lung fibroblasts (**Fig. 5B, S6B**).

**Figure 5.**
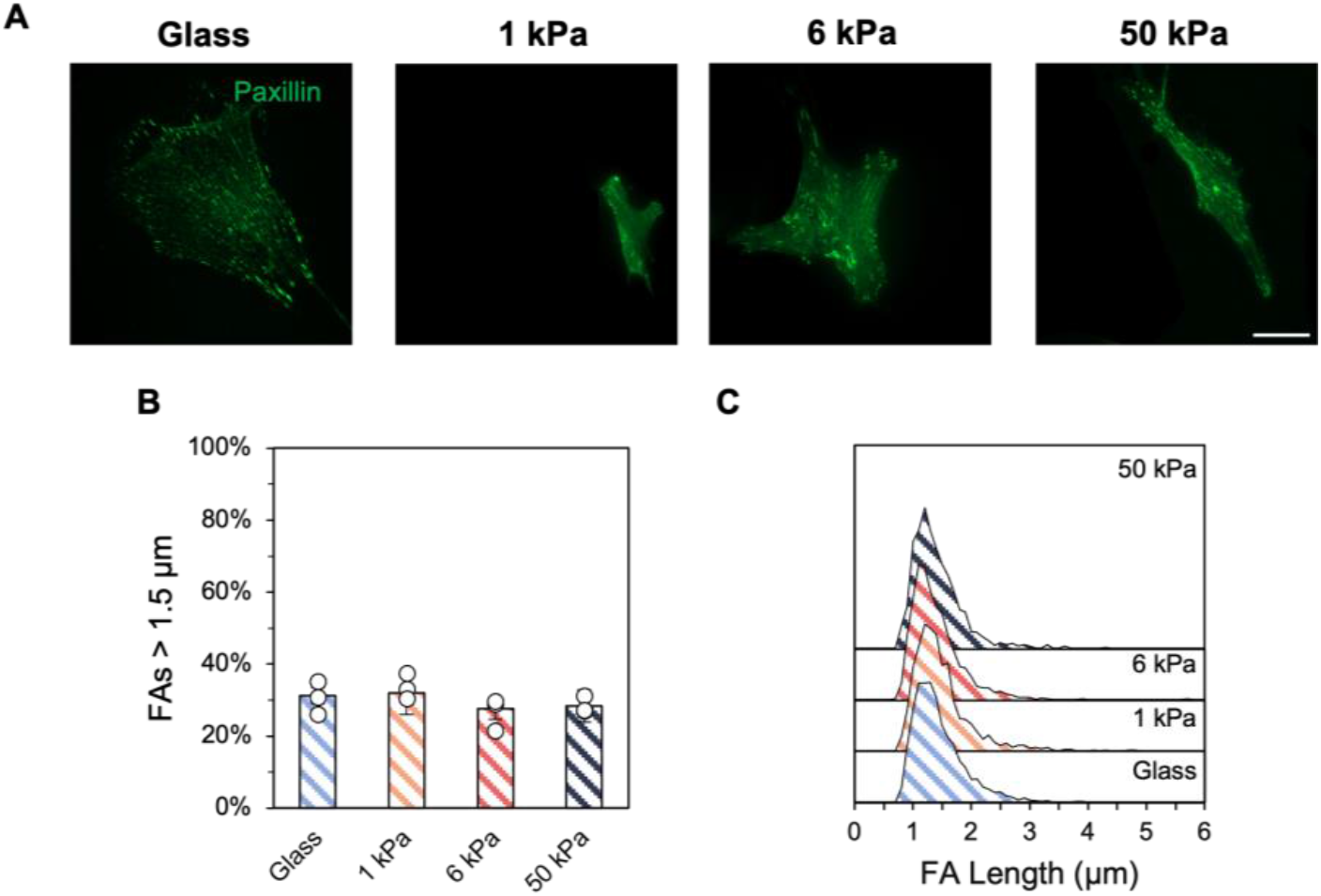
A) P5 hMSCs stained for paxillin to visualize focal adhesions, with quantification of B) focal adhesion lengths greater than 1.5 μm, a metric for mature adhesions, and C) ridgeline plots of the adhesion length distribution. Scale bar: 50 μm. *n* = 3 hydrogels per group. No statistically significant differences were observed.

When directly comparing cell spreading, roundness, MRTF-A nuclear localization, and focal adhesion size between early and late passage stromal cells, several important trends emerge (**Fig. 6, S7**). The first is that there are minimal differences in cell spread area and shape index (roundness) observed, although late passage lung fibroblasts are generally less round on compliant hydrogels (**Fig. S7B**). However, MRTF-A nuclear localization is generally lower for late passage hMSCs, with significant reductions for late passage hMSCs on hydrogels of 1 and 50 kPa compared to early passage hMSCs on the same stiffness substrates (**Fig. 6C**). Further, the number of mature (> 1.5 μm) focal adhesions was reduced for late passage hMSCs on stiffer hydrogels (**Fig. 6D**). Together, these results suggest a blunted response in later passage stromal cells to varying substrate stiffness.

**Figure 6.**
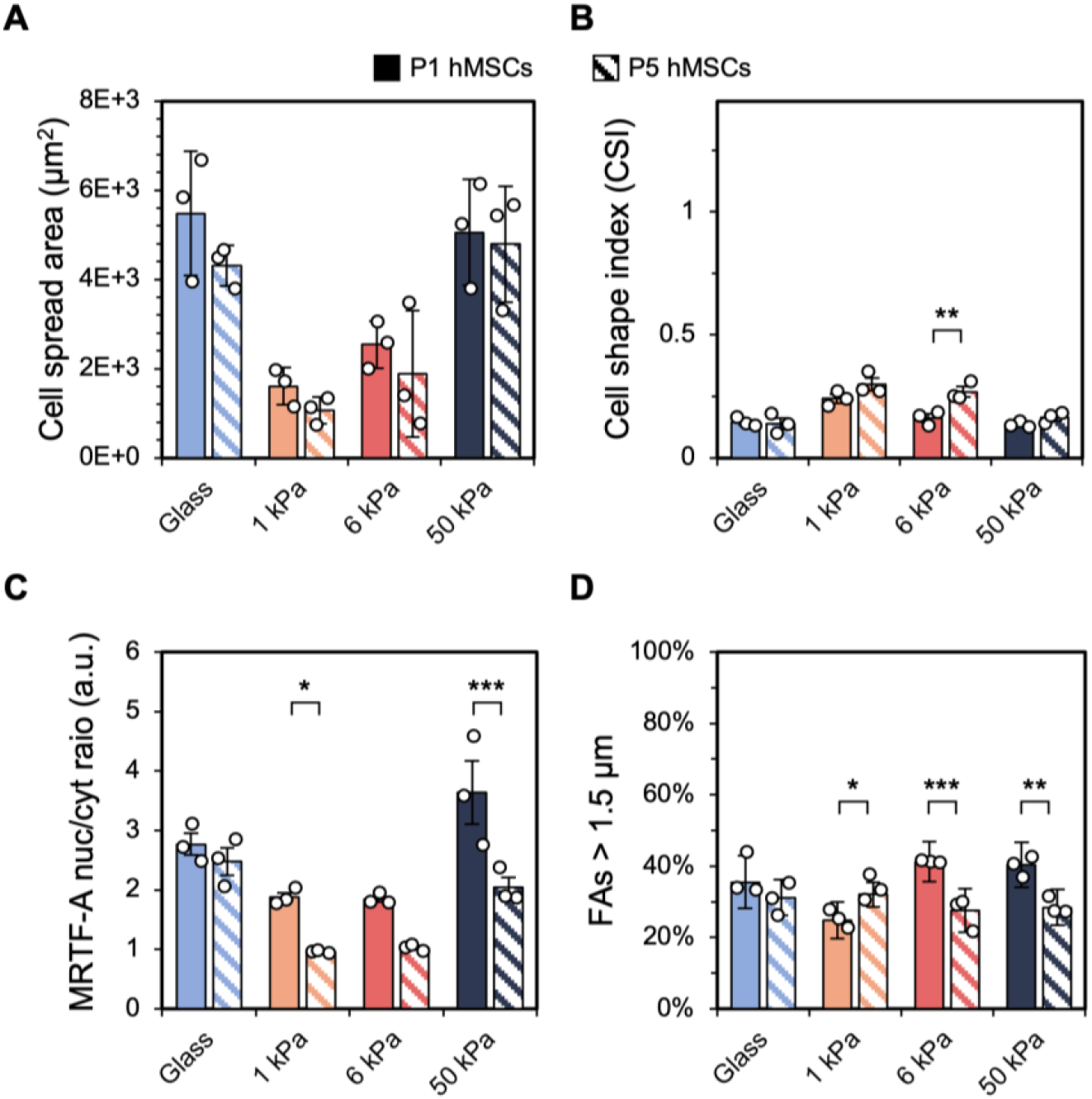
Comparison between P1 (solid bars) and P5 (striped bars) hMSCs with respect to A) cell spread area, B) cell shape index, C) MRTF-A nuclear*-to-* cytosol ratio, and D) focal adhesion lengths larger than 1.5 μm following 4 days of hydrogel culture. Data are reported as the mean ± S.E.M. *n* = 3 hydrogels per group. *** *P* <0.001, ** *P* < 0.01, * *P* < 0.05.

## 4. Discussion

While many studies have shown that cell behavior is influenced by environmental mechanical cues^15–19^, there is comparatively little information on how initial expansion on supraphysiologically stiff tissue culture plastic influences subsequent cell culture studies. Previous work on understanding mechanical memory highlighted the effects of serial passaging fibroblasts on top of silicone substrates, ranging from normal to fibrotic stiffnesses^38,39^, with sustained cell activation observed after mechanical priming on stiff substrates for 3 weeks^39^. Another study evaluated the effects of relatively short-term plastic priming (up to 10 days) on hMSC mechanosensitivity but did not evaluate the effects of serial passaging and TCP expansion^40^. Since many current research efforts are still performed using either primary or immortalized cells cultured extensively on TCP prior to *in vitro* hydrogel culture, it was of interest to investigate the influence of this approach on cell mechanosensitivity.

In the present study, the effect of hydrogel mechanical properties on the behavior of stromal cells that underwent variable levels of serial passaging was assessed. Cells were plated on either glass or hydrogels with stiffnesses of 1, 6, or 50 kPa mimicking normal or pathologic tissue stiffnesses. Both early and late passage hMSCs exhibited sensitivity to the mechanics of the underlying substrate during *in vitro* experiments as shown by distinct changes in spread area and formation of focal adhesions, as well as the nuclear localization of MRTF-A (**Fig. 2–5**). Similar observations have previously been made where increased stiffness drives greater cell spreading^17,50,51^, including for lung fibroblasts^39^, and increased focal adhesion maturation^52^. The correlation between increased spreading and increased MRTF-A nuclear localization has also been previously observed^36,53^. In comparing the behaviors of early versus late passage hMSCs, the late passage hMSCs exhibited a more blunted response to substrate stiffness as measured by these metrics, notably showing reduced MRTF-A nuclear localization and focal adhesion maturation (**Fig. 6**). Similar trends were observed when these same metrics were investigated using a fibroblast cell line immortalized by transfection with the catalytic subunit of human telomerase (hTERT) gene (**Fig. S3-S7**). Immortalized cell lines are manufactured to retain characteristic behaviors and can typically be passaged on TCP for much longer compared to their primary-derived counterparts, although phenotypic changes due to serial passaging are also known to occur. These findings highlight the implications of extended TCP expansion time on subsequent *in vitro* experiments to investigate both primary and immortalized cell behavior.

The effects of prolonged passaging on TCP has been evaluated for a variety of cell types, like primate brain microvessel endothelial cells, which displayed altered protein expression profiles and enzymatic activity^41^. In another study, primary porcine coronary arterial endothelial cells showed reduced proliferation, increased apoptosis, and p53 signaling activation, which facilitates tumor metastasis^42,43^. Furthermore, another study using human umbilical vein endothelial cells reported differences in cell spreading, shape, and migration with increased passaging^44^. In addition to primary cell cultures, these effects have also been observed for immortalized cell lines. For example, one study showed significant reductions in the expression of typical gene markers of uveal melanoma as a function of serial passaging^54^. This study also showed that serial passaging increased the tumorigenic potential of the cell line following subcutaneous injection into athymic mice^54^. Another report indicated that immortalized human colorectal adenocarcinoma cells showed variable morphological features with increased passaging as well as reduced growth kinetics and enzyme activity^55^. While those studies proved useful for understanding the roles that serial passaging can play on cell behavior, we wanted to specifically investigate changes in cellular mechanosensitivity. And, while our results may provide a level of understanding as to how prolonged TCP exposure affects subsequent cell behaviors, these effects will likely vary for different cell types and sources. Indeed, many studies have shifted toward culturing cells onto compliant substrates, rather than TCP, for a prolonged period to create conditions more closely resembling *in vivo* milieus^39,45,46^. One recent study showed that adaptation to TCP prior to the culture of a breast cancer cell line on polyacrylamide substrates resulted in reduced differences in morphology, including spread area, between cells on soft (1 kPa) or stiff (103 kPa) hydrogels^45^. In fact, a recent study showed that expanding cells on compliant surfaces could help erase this mechanical memory^38^ as a function of adaptation to the initially stiff environment of TCP, representing a potential future avenue of investigation.

Overall, this study evaluated how culture time influences subsequent stromal cell behavior in the context of TCP expansion prior to *in vitro* hydrogel culture. Our results underscore how initial passage length plays a role in cell response to physiologically relevant hydrogel models of disease with early passage stromal cells showing increased mechanosensitivity as measured by spreading, MRTF-A localization, focal adhesion organization, and chromatin condensation. Future experiments could consider extended culture times, different cell types/sources, and how more complex mechanical cues, like viscoelasticity or dynamic stiffening, influence the observed cell behaviors in this work.

## Supporting Information

^1^H NMR spectra for the hydrogel components, hydrogel mechanical characterization, and data from the fibroblast cell culture experiments can be found in the Supporting Information.

## Acknowledgments

This work was supported by the NSF (CAREER DMR/BMAT 2046592) and NIH (R35GM138187). The content is solely the responsibility of the authors and does not necessarily represent the official views of the National Institutes of Health.

## References

1 Wang, N., Tytell, J. D. & Ingber, D. E. Mechanotransduction at a distance: mechanically coupling the extracellular matrix with the nucleus. Nature Reviews Molecular Cell Biology 10, 75–82 (2009). https://doi.org:10.1038/nrm2594

2 Kolahi, K. S. & Mofrad, M. R. K. Mechanotransduction: a major regulator of homeostasis and development. Wiley Interdisciplinary Reviews: Systems Biology and Medicine 2, 625–639 (2010). https://doi.org:10.1002/wsbm.79

3 Wynn, T. A. Integrating mechanisms of pulmonary fibrosis. The Journal of Experimental Medicine 208, 1339–1350 (2011). https://doi.org:10.1084/jem.20110551

4 Wynn, T. A. Cellular and molecular mechanisms of fibrosis. J Pathol 214, 199–210 (2008). https://doi.org:10.1002/path.2277

5 Hinz, B. Mechanical Aspects of Lung Fibrosis. Proceedings of the American Thoracic Society 9, 137–147 (2012). https://doi.org:10.1513/pats.201202-017aw

6 Georges, P. C. et al. Increased stiffness of the rat liver precedes matrix deposition: implications for fibrosis. American Journal of Physiology-Gastrointestinal and Liver Physiology 293, G1147–G1154 (2007). https://doi.org:10.1152/ajpgi.00032.2007

7 Liu, F. et al. Feedback amplification of fibrosis through matrix stiffening and COX-2 suppression. The Journal of Cell Biology 190, 693–706 (2010). https://doi.org:10.1083/jcb.201004082

8 Parker, M. W. et al. Fibrotic extracellular matrix activates a profibrotic positive feedback loop. Journal of Clinical Investigation 124, 1622–1635 (2014). https://doi.org:10.1172/jci71386

9 Chen, H. et al. Mechanosensing by the α6-integrin confers an invasive fibroblast phenotype and mediates lung fibrosis. Nature Communications 7, 12564 (2016). https://doi.org:10.1038/ncomms12564

10 Liu, F. et al. Mechanosignaling through YAP and TAZ drives fibroblast activation and fibrosis. Am J Physiol Lung Cell Mol Physiol 308, L344–357 (2015). https://doi.org:10.1152/ajplung.00300.2014

11 Levental, K. R. et al. Matrix Crosslinking Forces Tumor Progression by Enhancing Integrin Signaling. Cell 139, 891–906 (2009). https://doi.org:10.1016/j.cell.2009.10.027

12 Calvo, F. et al. Mechanotransduction and YAP-dependent matrix remodelling is required for the generation and maintenance of cancer-associated fibroblasts. Nature Cell Biology 15, 637–646 (2013). https://doi.org:10.1038/ncb2756

13 Engler, A. J., Sen, S., Sweeney, H. L. & Discher, D. E. Matrix elasticity directs stem cell lineage specification. Cell 126, 677–689 (2006). https://doi.org:10.1016/j.cell.2006.06.044

14 Caliari, S. R. & Burdick, J. A. A practical guide to hydrogels for cell culture. Nat Methods 13, 405–414 (2016). https://doi.org:10.1038/nmeth.3839

15 Stowers, R. S., Allen, S. C. & Suggs, L. J. Dynamic phototuning of 3D hydrogel stiffness. Proceedings of the National Academy of Sciences 112, 1953–1958 (2015). https://doi.org:10.1073/pnas.1421897112

16 Blakney, A. K., Swartzlander, M. D. & Bryant, S. J. The effects of substrate stiffness on the in vitro activation of macrophages and in vivo host response to poly(ethylene glycol)-based hydrogels. Journal of Biomedical Materials Research Part A 100A, 1375–1386 (2012). https://doi.org:10.1002/jbm.a.34104

17 Guvendiren, M. & Burdick, J. A. Stiffening hydrogels to probe short- and long-term cellular responses to dynamic mechanics. Nature Communications 3, 792 (2012). https://doi.org:10.1038/ncomms1792

18 Evans, N. et al. Substrate stiffness affects early differentiation events in embryonic stem cells. European Cells and Materials 18, 1–14 (2009). https://doi.org:10.22203/ecm.v018a01

19 Caliari, S. R., Vega, S. L., Kwon, M., Soulas, E. M. & Burdick, J. A. Dimensionality and spreading influence MSC YAP/TAZ signaling in hydrogel environments. Biomaterials 103, 314–323 (2016). https://doi.org:10.1016/j.biomaterials.2016.06.061

20 Mabry, K. M., Lawrence, R. L. & Anseth, K. S. Dynamic stiffening of poly(ethylene glycol)-based hydrogels to direct valvular interstitial cell phenotype in a three-dimensional environment. Biomaterials 49, 47–56 (2015). https://doi.org:10.1016/j.biomaterials.2015.01.047

21 Chia, H. N., Vigen, M. & Kasko, A. M. Effect of substrate stiffness on pulmonary fibroblast activation by TGF-beta. Acta Biomater 8, 2602–2611 (2012). https://doi.org:10.1016/j.actbio.2012.03.027

22 Huang, X. et al. Matrix Stiffness-Induced Myofibroblast Differentiation Is Mediated by Intrinsic Mechanotransduction. American Journal of Respiratory Cell and Molecular Biology 47, 340–348 (2012). https://doi.org:10.1165/rcmb.2012-0050oc

23 Caliari, S. R. et al. Stiffening hydrogels for investigating the dynamics of hepatic stellate cell mechanotransduction during myofibroblast activation. Sci Rep 6, 21387 (2016). https://doi.org:10.1038/srep21387

24 Duscher, D. et al. Mechanotransduction and fibrosis. Journal of Biomechanics 47, 1997–2005 (2014). https://doi.org:10.1016/jjbiomech.2014.03.031

25 Humphrey, J. D., Dufresne, E. R. & Schwartz, M. A. Mechanotransduction and extracellular matrix homeostasis. Nature Reviews Molecular Cell Biology 15, 802–812 (2014). https://doi.org:10.1038/nrm3896

26 Ingber, D. E. Cellular mechanotransduction: putting all the pieces together again. FASEB J 20, 811–827 (2006). https://doi.org:10.1096/fj.05-5424rev

27 Seong, J., Wang, N. & Wang, Y. Mechanotransduction at focal adhesions: from physiology to cancer development. J Cell Mol Med 17, 597–604 (2013). https://doi.org:10.1111/jcmm.12045

28 Jansen, K. A., Atherton, P. & Ballestrem, C. Mechanotransduction at the cell-matrix interface. Seminars in Cell & Developmental Biology 71, 75–83 (2017). https://doi.org:10.1016/j.semcdb.2017.07.027

29 Parsons, J. T., Horwitz, A. R. & Schwartz, M. A. Cell adhesion: integrating cytoskeletal dynamics and cellular tension. Nature Reviews Molecular Cell Biology 11, 633–643 (2010). https://doi.org:10.1038/nrm2957

30 Gardel, M. L., Schneider, I. C., Aratyn-Schaus, Yvonne & Waterman, C. M. Mechanical Integration of Actin and Adhesion Dynamics in Cell Migration. Annual Review of Cell and Developmental Biology 26, 315–333 (2010). https://doi.org:10.1146/annurev.cellbio.011209.122036

31 Zimerman, B., Volberg, T. & Geiger, B. Early molecular events in the assembly of the focal adhesion-stress fiber complex during fibroblast spreading. Cell Motility and the Cytoskeleton 58, 143–159 (2004). https://doi.org:10.1002/cm.20005

32 Harjanto, D. & Zaman, M. H. Matrix mechanics and receptor–ligand interactions in cell adhesion. Org. Biomol. Chem. 8, 299–304 (2010). https://doi.org:10.1039/b913064k

33 Kishi, T., Mayanagi, T., Iwabuchi, S., Akasaka, T. & Sobue, K. Myocardin-related transcription factor A (MRTF-A) activity-dependent cell adhesion is correlated to focal adhesion kinase (FAK) activity. Oncotarget 7, 72113–72130 (2016). https://doi.org:10.18632/oncotarget.12350

34 Parmacek, M. S. Myocardin-Related Transcription Factors. Circulation Research 100, 633–644 (2007). https://doi.org:10.1161/01.res.0000259563.61091.e8

35 Zhao, X. H. et al. Force activates smooth muscle alpha-actin promoter activity through the Rho signaling pathway. J Cell Sci 120, 1801–1809 (2007). https://doi.org:10.1242/jcs.001586

36 O’Connor, J. W. & Gomez, E. W. Cell adhesion and shape regulate TGF-beta1-induced epithelial-myofibroblast transition via MRTF-A signaling. PLoS One 8, e83188 (2013). https://doi.org:10.1371/journal.pone.0083188

37 Hinson, J. S., Medlin, M. D., Lockman, K., Taylor, J. M. & Mack, C. P. Smooth muscle cell-specific transcription is regulated by nuclear localization of the myocardin-related transcription factors. Am J Physiol Heart Circ Physiol 292, H1170–1180 (2007). https://doi.org:10.1152/ajpheart.00864.2006

38 Li, C. X. et al. MicroRNA-21 preserves the fibrotic mechanical memory of mesenchymal stem cells. Nat Mater 16, 379–389 (2017). https://doi.org:10.1038/nmat4780

39 Balestrini, J. L., Chaudhry, S., Sarrazy, V., Koehler, A. & Hinz, B. The mechanical memory of lung myofibroblasts. Integr Biol (Camb) 4, 410–421 (2012). https://doi.org:10.1039/c2ib00149g

40 Yang, C., Tibbitt, M. W., Basta, L. & Anseth, K. S. Mechanical memory and dosing influence stem cell fate. Nat Mater 13, 645-652 (2014). https://doi.org:10.1038/nmat3889

41 Shi, Q., Aida, K., Vandeberg, J. L. & Wang, X. L. Passage-Dependent Changes in Baboon Endothelial Cells—Relevance to *In Vitro* Aging. DNA and Cell Biology 23, 502–509 (2004). https://doi.org:10.1089/1044549041562294

42 Lee, M. Y. K., Sørensen, G. L., Holmskov, U. & Vanhoutte, P. M. The presence and activity of SP-D in porcine coronary endothelial cells depend on Akt/PI3K, Erk and nitric oxide and decrease after multiple passaging. Molecular Immunology 46, 1050–1057 (2009). https://doi.org:10.1016/j.molimm.2008.09.027

43 Lee, M. Y. K., Wang, Y. & Vanhoutte, P. M. Senescence of Cultured Porcine Coronary Arterial Endothelial Cells Is Associated with Accelerated Oxidative Stress and Activation of NFκB. Journal of Vascular Research 47, 287–298 (2010). https://doi.org:10.1159/000265563

44 Liao, H. et al. Effects of long-term serial cell passaging on cell spreading, migration, and cell-surface ultrastructures of cultured vascular endothelial cells. Cytotechnology 66, 229–238 (2014). https://doi.org:10.1007/s10616-013-9560-8

45 Syed, S., Schober, J., Blanco, A. & Zustiak, S. P. Morphological adaptations in breast cancer cells as a function of prolonged passaging on compliant substrates. PLOS ONE 12, e0187853 (2017). https://doi.org:10.1371/journal.pone.0187853

46 Cabriales, L. et al. Hepatic C9 cells switch their behaviour in short or long exposure to soft substrates. Biology of the Cell 112, 265–279 (2020). https://doi.org:10.1111/boc.201900115

47 Rodriguez, N. M., Desai, R. A., Trappmann, B., Baker, B. M. & Chen, C. S. Micropatterned multicolor dynamically adhesive substrates to control cell adhesion and multicellular organization. Langmuir 30, 1327–1335 (2014). https://doi.org:10.1021/la404037s

48 Berginski, M. E. & Gomez, S. M. The Focal Adhesion Analysis Server: a web tool for analyzing focal adhesion dynamics. F1000Research 2, 68 (2013). https://doi.org:10.12688/f1000research.2-68.v1

49 Booth, A. J. et al. Acellular normal and fibrotic human lung matrices as a culture system for in vitro investigation. Am J Respir Crit Care Med 186, 866–876 (2012). https://doi.org:10.1164/rccm.201204-0754OC

50 Olsen, A. L. et al. Hepatic stellate cells require a stiff environment for myofibroblastic differentiation. Am J Physiol Gastrointest Liver Physiol 301, G110–118 (2011). https://doi.org:10.1152/ajpgi.00412.2010

51 Yeh, Y.-C. et al. Mechanically dynamic PDMS substrates to investigate changing cell environments. Biomaterials 145, 23–32 (2017). https://doi.org:10.1016/j.biomaterials.2017.08.033

52 Hui, E., Moretti, L., Barker, T. H. & Caliari, S. R. The combined influence of viscoelasticity and adhesive cues on fibroblast spreading and focal adhesion formation Cell Mol Bioeng 14, 427–440 (2021).

53 Hui, E., Gimeno, K. I., Guan, G. & Caliari, S. R. Spatiotemporal control of viscoelasticity in phototunable hyaluronic acid hydrogels. Biomacromolecules 20, 4126–4134 (2019). https://doi.org:10.1021/acs.biomac.9b00965

54 Mouriaux, F. et al. Effects of Long-term Serial Passaging on the Characteristics and Properties of Cell Lines Derived From Uveal Melanoma Primary Tumors. Invest Ophthalmol Vis Sci 57, 5288–5301 (2016). https://doi.org:10.1167/iovs.16-19317

55 Briske-Anderson, M. J., Finley, J. W. & Newman, S. M. The influence of culture time and passage number on the morphological and physiological development of Caco-2 cells. Proc Soc Exp Biol Med 214, 248–257 (1997). https://doi.org:10.3181/00379727-214-44093

