## Supporting Information for "Serial passaging affects stromal cell mechanosensitivity on hyaluronic acid hydrogels"

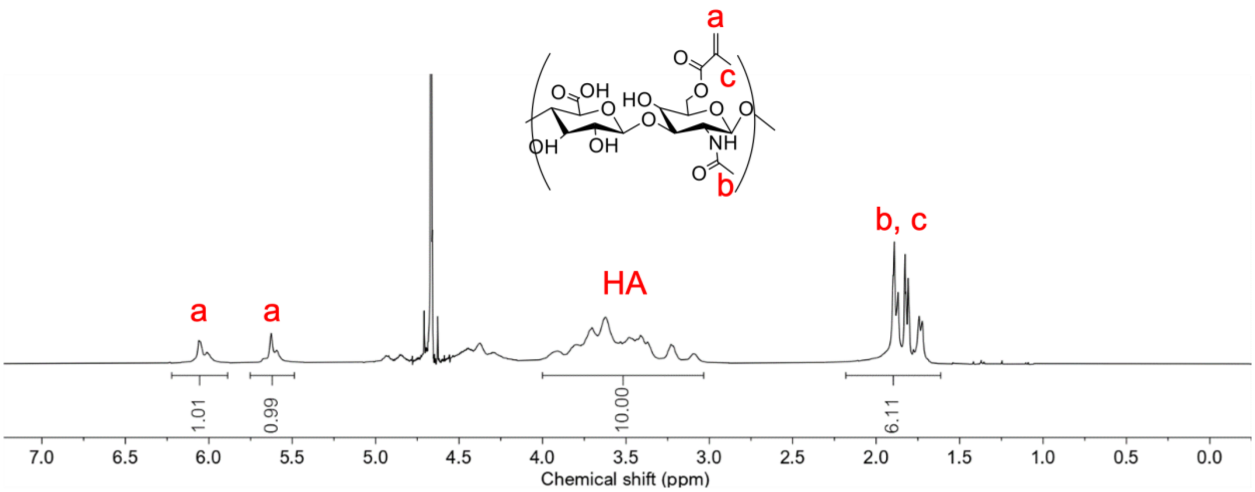


**Figure S1**. $^{1}H$ NMR spectrum of methacrylated hyaluronic acid (MeHA). The degree of modification was determined to be 100% as determined by the alkene peaks labeled a.

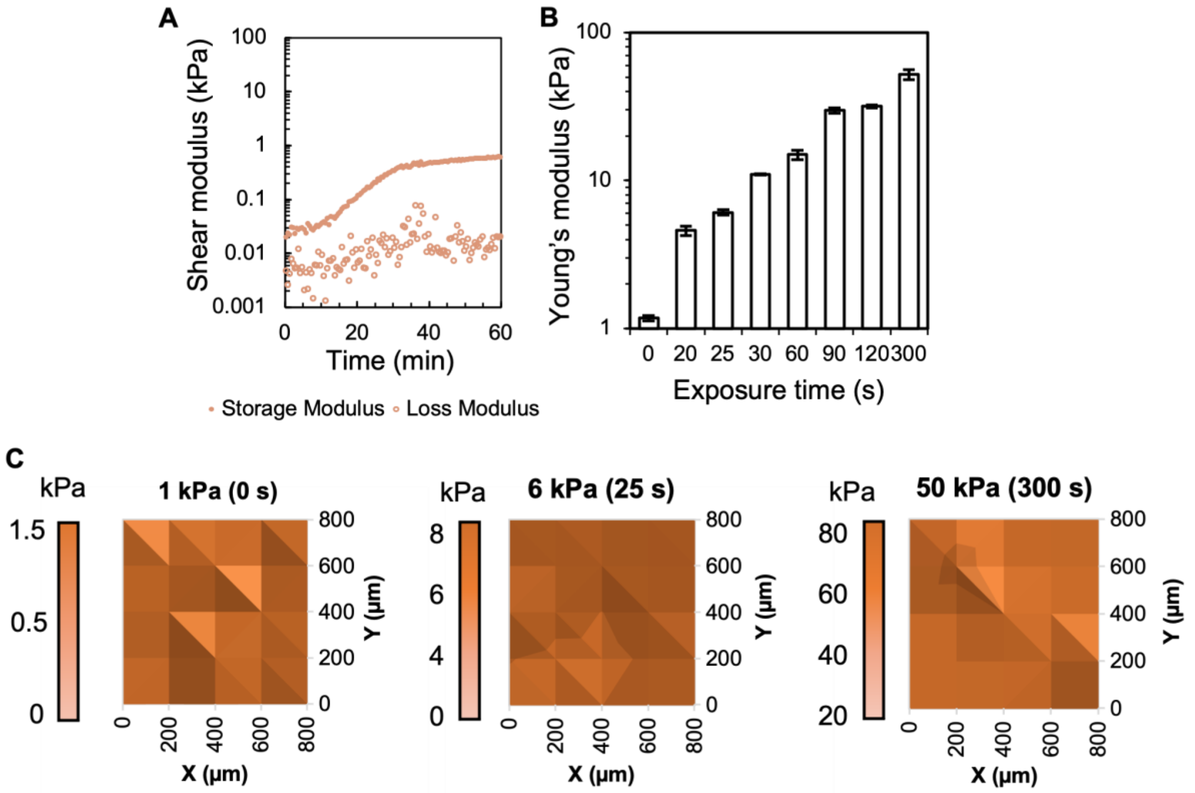


**Figure S2**. Mechanical characterization of MeHA hydrogels. **A**) *In situ* rheology of initial network formation. **B**) Nanoindentation of hydrogels following increasing lengths of blue light exposure in the presence of 2.2 mM LAP photoinitiator. **C**) Surface maps from nanoindentation to illustrate mechanical homogeneity across the hydrogels.

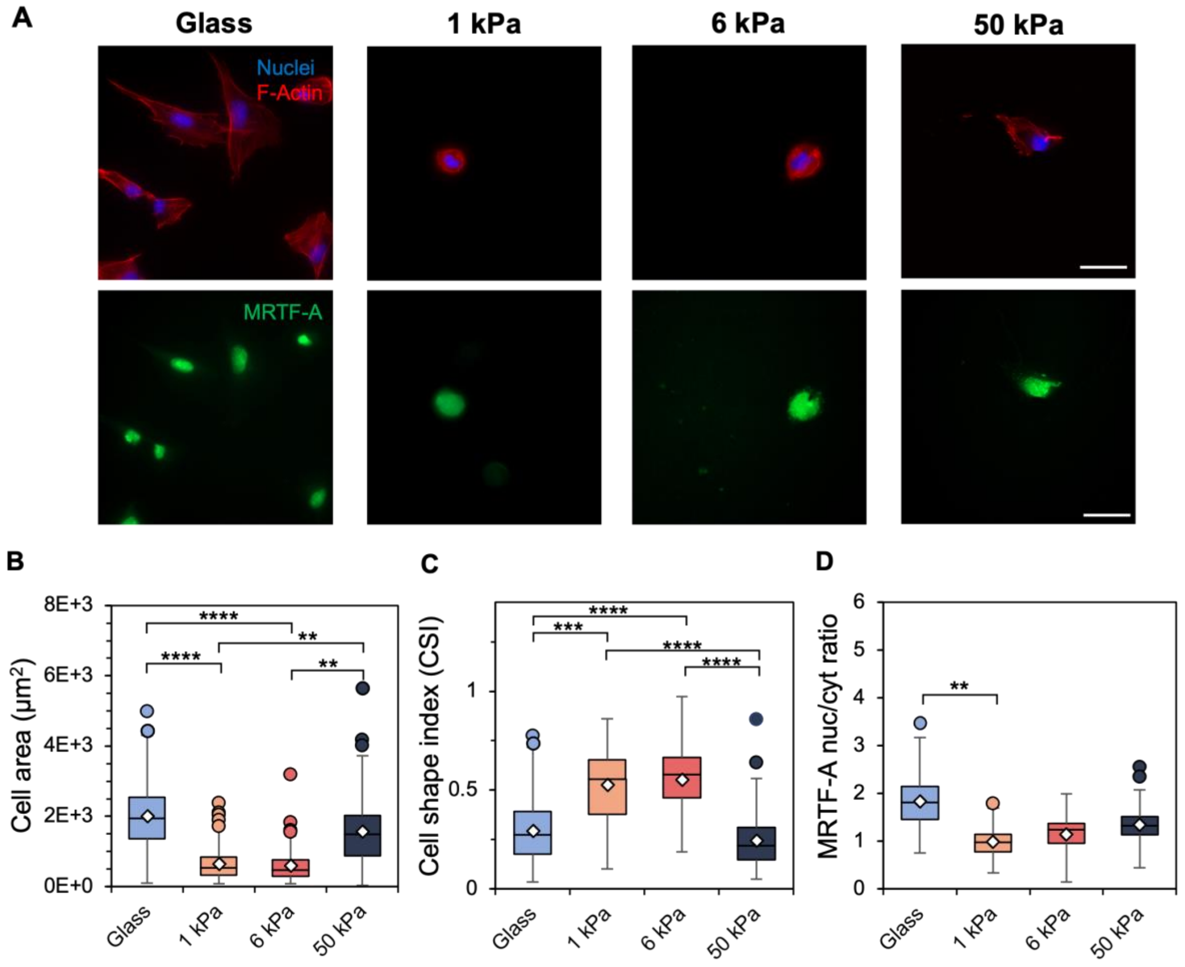


**Figure S3**. **A**) Representative images of passage 1 (P1) fibroblasts cultured on glass and 1, 6, and 50 kPa hydrogels for 4 days. Scale bars: 50 µm. Following 4 days of culture, quantification of **B**) cell spread area (µm$^2$), **C**) cell shape index, a measure of cell circularity, and **D**) MRTF-A nuclear-to-cytosol ratio was performed. $n$ = 3 hydrogels per group. **** $P < 0.0001$, *** $P < 0.001$, ** $P < 0.01$.

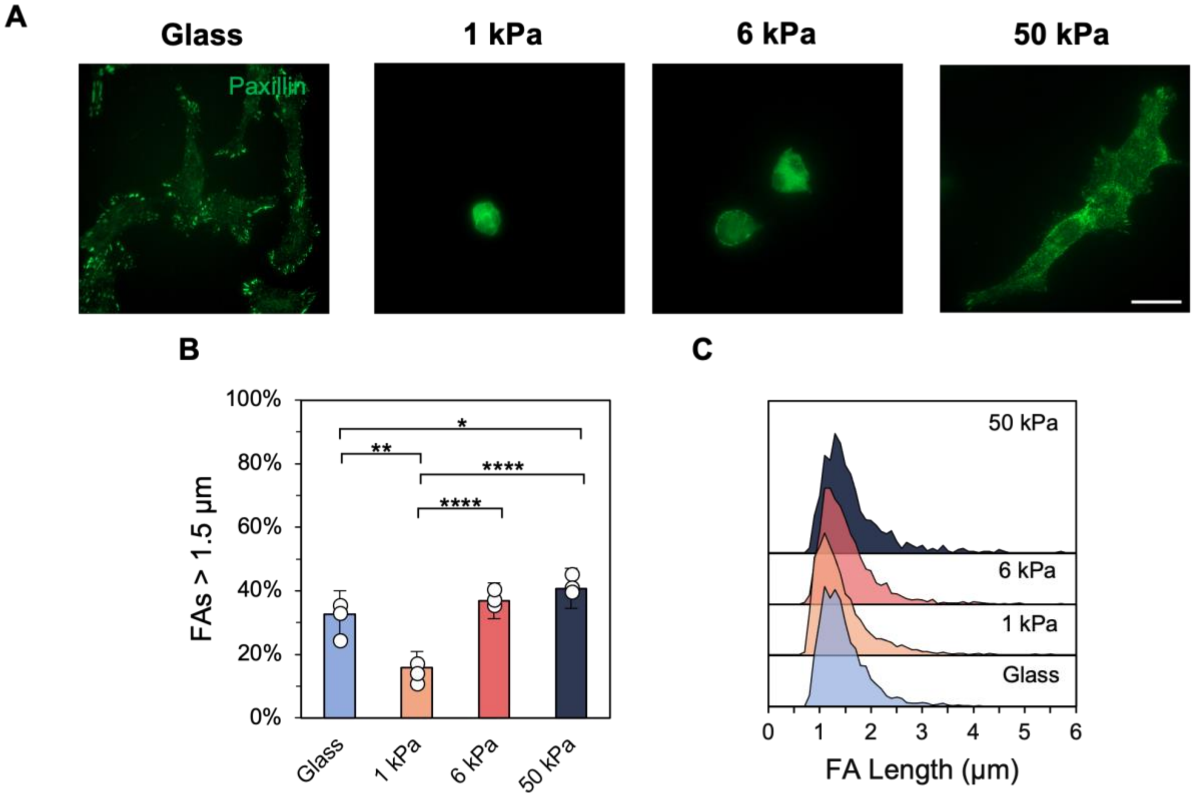


**Figure S4**. **A**) P1 fibroblasts were stained for paxillin to visualize focal adhesions, with quantification of **B**) focal adhesion lengths larger than 1.5 μm, a metric for mature adhesions, and **C**) ridgeline plots of the adhesion length distribution. Scale bar: 50 μm. $n = 3$ hydrogels per group. **** $P < 0.0001$, ** $P < 0.01$, * $P < 0.05$.

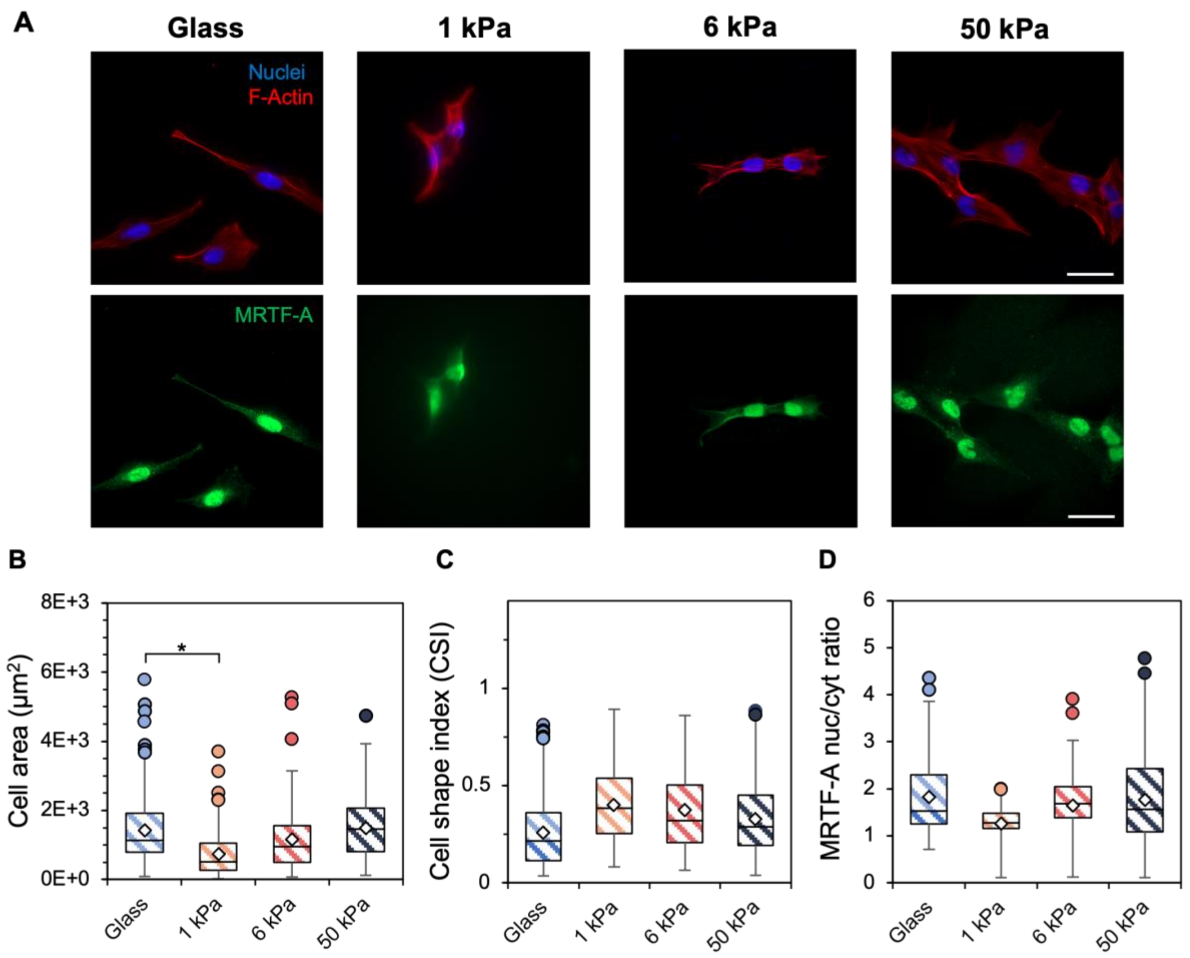


**Figure S5**. **A**) Representative images of passage 10 (P10) fibroblasts cultured on glass and 1, 6, and 50 kPa hydrogels for 4 days. Scale bars: 50 µm. Following 4 days of culture, quantification of **B**) cell spread area ($\mu m^2$), **C**) cell shape index, a measure of cell circularity, and **D**) MRTF-A nuclear-to-cytosol ratio was performed. $n$ = 3 hydrogels per group. * $P < 0.05$.

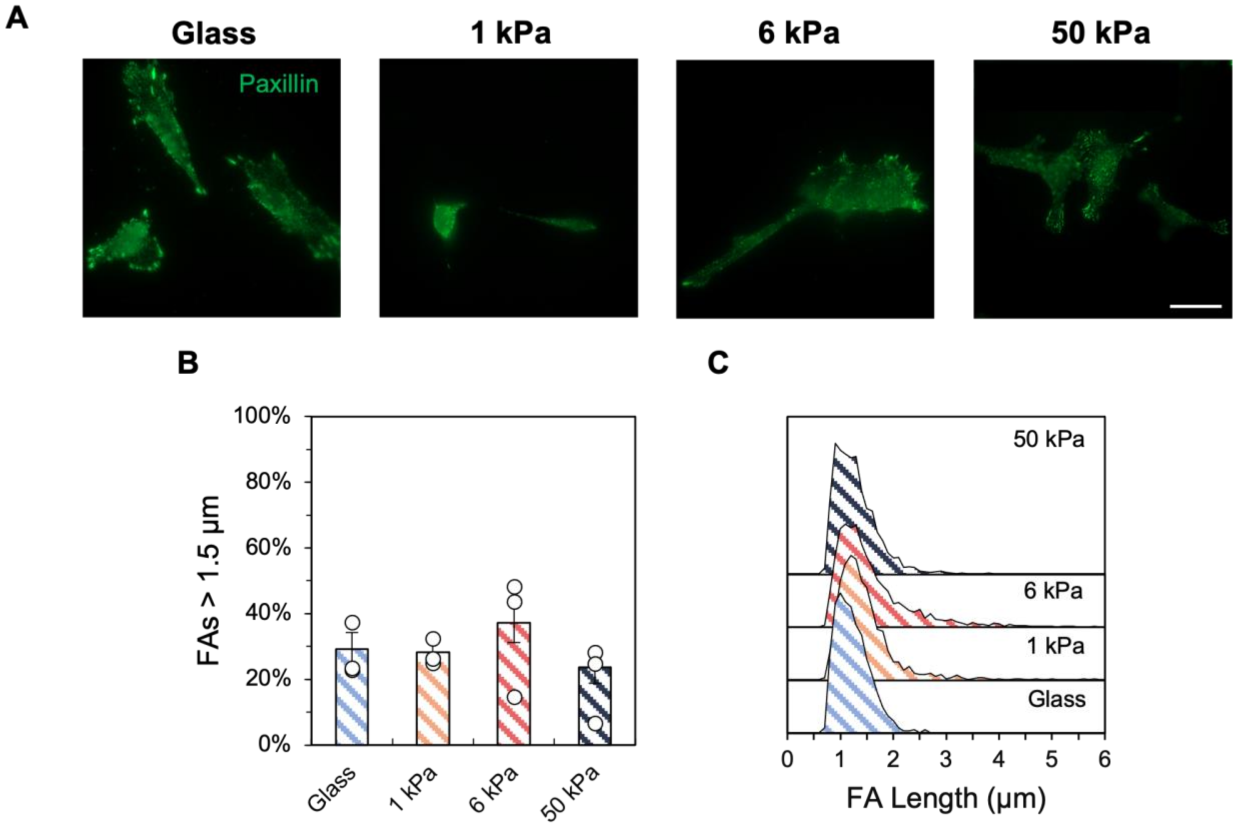


**Figure S6**. **A**) P10 fibroblasts were stained for paxillin to visualize focal adhesions, with quantification of **B**) focal adhesion lengths larger than 1.5 µm, a metric for mature adhesions, and **C**) ridgeline plots of the adhesion length distribution. Scale bar: 50 µm. $n$ = 3 hydrogels per group. No statistically significant differences were observed.

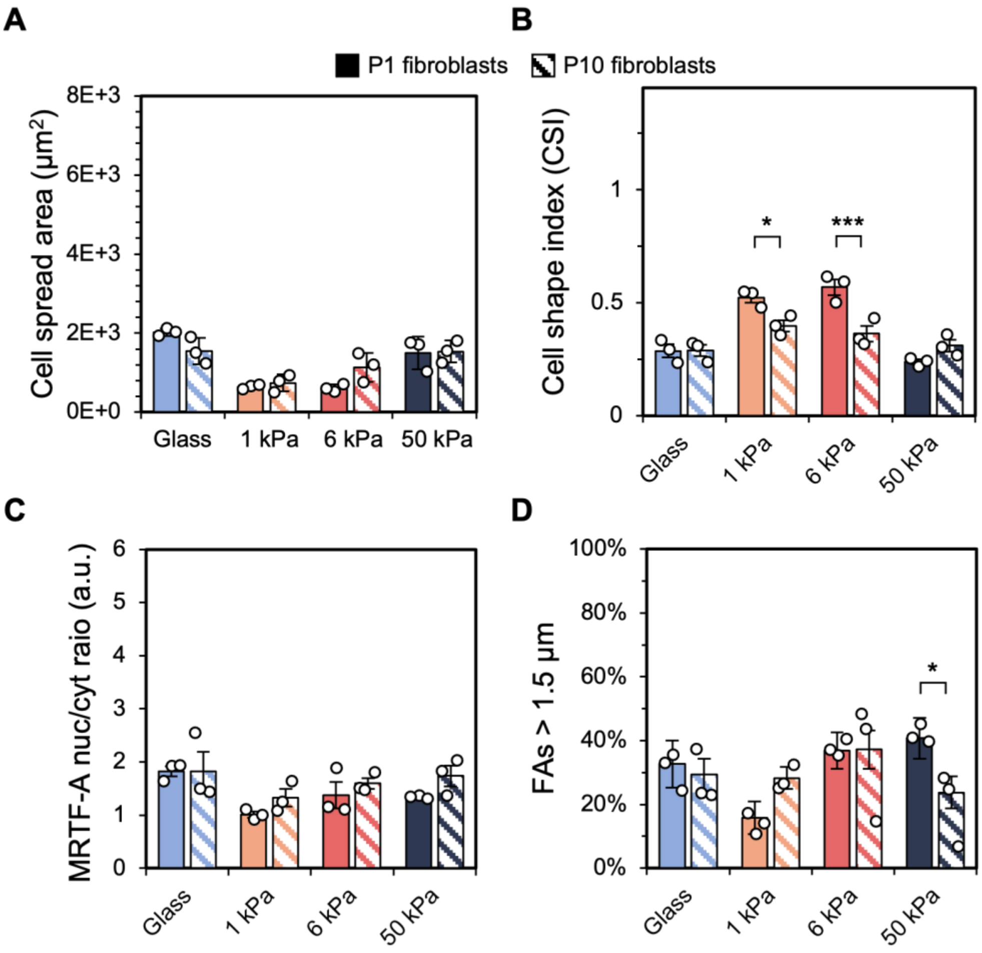


**Figure S7**. Comparison between P1 (solid bars) and P10 (striped bars) fibroblasts with respect to **A**) cell spread area, **B**) cell shape index, **C**) MRTF-A nuclear-to-cytosol ratio, and **D**) focal adhesion lengths larger than 1.5 µm following 4 days of hydrogel culture. Data are reported as the mean ± S.E.M. $n = 3$ hydrogels per group. * $P < 0.05$.
